# Limited consequences for loss of RNA-directed DNA methylation in *Setaria viridis* domains rearranged methyltransferase (DRM) mutants

**DOI:** 10.1101/2022.01.05.474142

**Authors:** Andrew Read, Trevor Weiss, Peter A Crisp, Zhikai Liang, Jaclyn Noshay, Claire C Menard, Chunfang Wang, Meredith Song, Candice N Hirsch, Nathan M Springer, Feng Zhang

## Abstract

The Domains Rearranged Methyltransferases (DRMs) are crucial for RNA-directed DNA methylation (RdDM) in plant species. *Setaria viridis* is a model monocot species with a relatively compact genome that has limited transposable element content. CRISPR-based genome editing approaches were used to create loss-of-function alleles for the two putative functional DRM genes in *S. viridis* to probe the role of RdDM. The analysis of *drm1ab* double mutant plants revealed limited morphological consequences for the loss of RdDM. Whole-genome methylation profiling provided evidence for wide-spread loss of methylation in CHH sequence contexts, particularly in regions with high CHH methylation in wild-type plants. Evidence was also found for locus-specific loss of CG and CHG methylation, even in some regions that lack CHH methylation. Transcriptome profiling identified a limited number of genes with altered expression in the *drm1ab* mutants. The majority of genes with elevated CHH methylation directly surrounding the transcription start site or in nearby promoter regions do not have altered expression in the *drm1ab* mutant even when this methylation is lost, suggesting limited regulation of gene expression by RdDM. Detailed analysis of the expression of transposable elements identified several transposons that are transcriptionally activated in *drm1ab* mutants. These transposons likely require active RdDM for maintenance of transcriptional repression.

**Significance statement:** Methylation profiling of *Setaria viridis* plants that lack functional Domains Rearranged Methyltransferase genes reveal widespread loss of DNA methylation in the CHH sequence context. Transcriptome analysis reveals a small set of genes and transposons that are silenced by RNA-directed DNA methylation.

## Introduction

DNA methylation is a common chromatin modification in many plant genomes. Cytosine methylation is the result of post-replication modification that adds a methyl group to the 5’ carbon. While virtually all plants that have been assessed contain DNA methylation, there are differences in the levels and context-specific patterns of methylation in different species (Niederhuth *et al*., 2016). The majority of our knowledge about the molecular mechanisms that control DNA methylation and the functions of DNA methylation are based on studies in *Arabidopsis thaliana* (Arabidopsis) due to the viability of plants with highly reduced DNA methylation (Law and Jacobsen, 2010; Matzke and Mosher, 2014). However, studies in other plants have suggested differences in the patterns and control of DNA methylation (Springer et al. 2016; Niederhuth et al. 2016).

DNA methylation in plant genomes involves several distinct methyltransferases that create or maintain DNA methylation and these can be distinguished by the local sequence context (Law and Jacobsen, 2010). CG methylation is often present at high levels in plant genomes and is maintained following DNA replication due to the preference of MET1 and orthologous genes for hemimethylated sites (Law and Jacobsen, 2010). CHG (H = A, T or C) methylation is also quite common and is catalyzed by chromomethylase enzymes in a feed-forward loop with H3K9me2 (Du *et al*., 2012; Johnson *et al*., 2007). CHH methylation occurs at non-symmetrical genomic sites and requires specific targeting mechanisms. Evidence from Arabidopsis suggests two distinct pathways to maintain or create CHH methylation (Zemach *et al*., 2013; Stroud *et al*., 2014; Cao and Jacobsen, 2002b). The RNA-directed DNA methylation (RdDM) pathway utilizes PolIV and PolV to generate and utilize 24nt sRNAs to target the DRM genes to specific loci (Matzke and Mosher, 2014). Most of the RdDM activity is focused on either small TEs or the edges of longer TEs (Zemach *et al*., 2013). In maize, the RdDM activity seems to be particularly high at the edges of TEs near expressed genes (Li *et al*., 2015; Gent *et al*., 2013). In Arabidopsis, the CHH methylation found within internal regions of longer TEs requires activity of the CMT2 chromomethylase (Zemach *et al*., 2013; Stroud *et al*., 2014).

The *DOMAINS REARRANGED METHYLTRANSFERASES (DRM)* genes were identified as putative relatives of the mammalian *de novo* methyltransferase *Dnmt3* with a unique rearrangement for the order of the methyltransferase domains (Cao *et al*., 2000). Studies in Arabidopsis indicated the *drm1/drm2* mutants had reduced CHH methylation and were compromised for silencing of some genes and TEs (Cao *et al*., 2003; Tran *et al*., 2005; Stroud, Greenberg, *et al*., 2013; Chan *et al*., 2004; Cao and Jacobsen, 2002a). However, there are no substantial developmental or morphological abnormalities in Arabidopsis plants that lack *DRM1/2* (Cao and Jacobsen, 2002a; Chan *et al*., 2006). Combining the *drm* mutant with a loss of function for *CHROMOMETHYLASE3 (CMT3)* results in significant phenotypic impacts suggesting partially redundant control of gene silencing and asymmetric methylation by *DRM* and *CMT* genes (Chan *et al*., 2006; Henderson and Jacobsen, 2008; Stroud *et al*., 2014). In Arabidopsis *drm1/drm2* mutants there are substantial reductions of CHH methylation at many loci and there is partial reduction of CHG at these same regions that is completely reduced in *drm1 drm2 cmt3* mutants suggesting combined control of CHH and CHG methylation by DRM and CMT at these sites (Stroud *et al*., 2014).

CHH methylation triggered by RNA directed DNA methylation (RdDM) appears to play a limited role in regulating expression (Cao and Jacobsen, 2002b; Stroud *et al*., 2014). There are certainly some endogenous loci and transgenes that are silenced by DRM or other components of the RdDM machinery. However, the number of genes or transposons that are activated in a *drm1/drm2* mutant is limited, and it seems that there is substantial redundancy between *DRM* and *CMT* pathways to maintain silencing (Stroud *et al*., 2014). Loss-of-function for the rice *DRM* ortholog *OsDRM2* results in pleiotropic phenotypes as well as aberrant expression of some transposons and genes (Moritoh *et al*., 2012; Tan *et al*., 2016).

In this study we use CRISPR to create loss-of-function alleles for the two putative functional DRM orthologs in *Setaria viridis. S. viridis* is a model C4 grass that has a relatively small genome and a short generation time (Brutnell *et al*., 2010; Mamidi *et al*., 2020; Thielen *et al*., 2020; Bennetzen *et al*., 2012) and may provide a model system for analysis of the control of DNA methylation in monocots. A high quality reference genome is available for the A10 *S. viridis* accession (Mamidi et al. 2020) and there is also a reference genome for the highly transformable accession ME034V (Thielen et al. 2020). We find that loss of *DRM* results in substantial reductions in CHH methylation. There are also losses of CHG and CG methylation at regions with reduced CHH methylation as well as within regions that lack CHH methylation in both wild-type and mutant plants. Despite the widespread changes in CHH methylation there are limited changes in gene expression in plants lacking functional *DRM* genes. A subset of transposons are transcriptionally activated in the *DRM* mutant lines, highlighting the role of RdDM in regulation of transposons.

## Results

### Isolation of DRM loss-of-function alleles

The *Setaria viridis* genome has two genes that encode putatively functional *DRM* genes; *Drm1a* - Sevir.9G574800 (A10) / Svm9G0069770 (ME034V) and *Drm1b* - Sevir.9G496200 (A10) / Svm9G0060600 (ME034V). These two genes are located ~5 Mb apart on chromosome 9 and likely represent a relatively recent duplication event (Figure S1A). The two protein sequences have 67.1% identity and 76.5% similarity. Expression atlas data for both accession A10 and ME034v suggest that *SvDrm1a* is much more highly expressed than *Drm1b* in leaf tissue (Figure S1B-D). This suggests that *Drm1a* may have more functional relevance but there is potential redundancy for these two genes. There is also a *DRM3*-like gene, Sevir.3G052500 (A10) / Svm3G0006370 (ME034v), that lacks critical residues in the catalytic domain and is unlikely to provide functional methyltransferase activity. This is likely an ortholog of the Arabidopsis *DRM3*, which encodes a catalytically inert protein that appears to be required as a cofactor for proper CHH methylation at some loci (Henderson *et al*., 2010; Costa-Nunes *et al*., 2014)

A total of three guide RNAs (gRNAs) targeting *Drm1a* and *Drm1b* were designed as described in our previous study (Weiss et al. 2020). To generate *S. viridis* mutant plants with double gene knockouts, the T-DNA construct (pTW45) expressing Cas9_Trex2 with all three gRNA sequences was transformed through agrobacterium-mediated transformation into the transformable *S. viridis* genotype ME034V (Weiss et al. 2020) (Figure S2A;_see methods for details). T0 plants with edits at both targeted genes were identified and selected for self-pollination. Progeny were screened as in Weiss et al 2020, and two transgene free T1 plants were selected for further propagation: one containing edits at *drm1a* and *drm1b* (12-9), and one plant with WT alleles (84-27). To identify homozygous progenies with frame-shift mutations at both genes, two generations of self pollination were performed. T3 plants with the edits at both genes were identified (hereafter referred to as *drm1ab*), which included a 3 bp deletion in *Drm1a* that introduces an early stop codon as well as a 2bp and a 6bp deletion in *Drm1b* that results in a frameshift mutation (Figure 1A, Figure S2B). The predicted proteins produced by these mutant alleles both lack critical domains that are necessary for methyltransferase activity. The *drm1ab* plants are fully viable (Figure 1B). The *drm1ab* plants are reduced in stature and have reduced leaf length (Figure 1B-C). In addition, the *drm1ab* plants exhibit delayed flowering relative to wild-type. The severity of the change in stature and flowering time were variable in different growth conditions. While the *drm1ab* plants exhibit reduced stature, there were no major morphological differences noted in vegetative or floral architecture.

**Figure 1.**
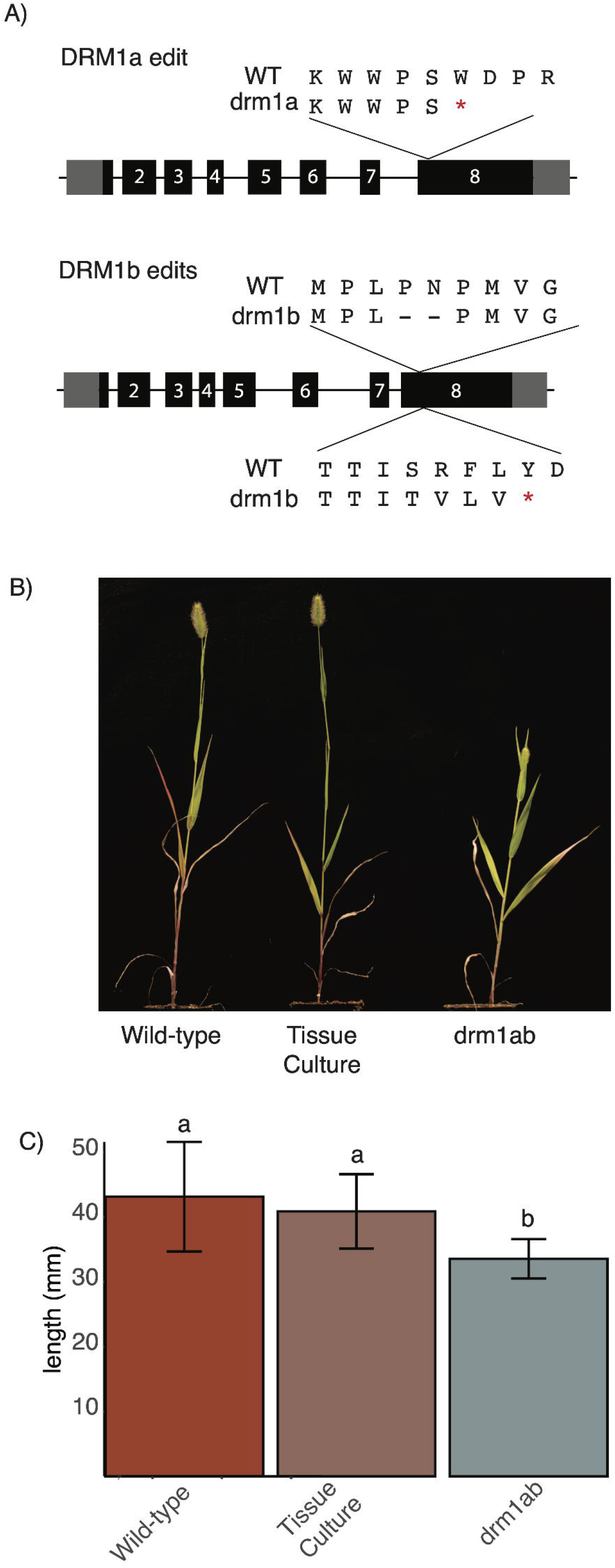
Isolation of loss-of-function alleles for DRM genes in *Setaria viridis*. (A) Sequencing of transgene-free plants derived from transgenic parents expressing gRNAs targeting the *Drm1a* and *Drm1b* gene identified individuals that are homozygous for mutations at both target genes. The schematic indicates the position and sequence change at each locus. (B) Images showing wild-type ME034V and *drm1ab* double mutant plants. (C) Leaf length is reduced in the *drm1ab* mutant plants relative to wild-type controls or the progeny of plants derived from tissue-culture. Letters indicate significant differences (p<.05).

### Characterization of methylation domains within the Setaria viridis genome

Whole genome DNA methylation profiles were generated for a single replicate sample of wild-type *S. viridis* ME034V as well as three biological replicates from plants whose parent (84-27) was regenerated from tissue culture and three biological replicates of transgene-free *drm1ab* plants. All samples were collected from seedling leaf tissue at a developmental stage in which there are no phenotypic differences between the mutant and wild-type plants. Enzymatic conversion rates for all samples ranged from 99.43 to 99.81%. The genome-wide DNA methylation levels for wild-type ME034V plants (Figure 2A, S3A) are quite similar to reported levels for *S. viridis* accession, A10 (Niederhuth *et al*., 2016). Prior studies have suggested some changes in DNA methylation induced due to tissue culture in rice and maize (Stroud, Ding, *et al*., 2013; Han *et al*., 2018). We did not observe significant differences in the overall DNA methylation levels between wild-type and tissue-culture derived samples (Figure S3A). The genome was divided into 100 bp tiles, each of which was classified based on the levels of CG, CHG and CHH methylation (Figure S3B; see Methods for details). The wild-type and tissue culture derived plants had very similar proportions of the genome classified as high CG and CHG (~28% of genome), CG-only (~15% of genome), or high CHH (~1.4% of genome) (Figure S3B). For the analysis of *drm1ab* we focused on contrasts between the three biological replicates of tissue-culture derived plants and the loss-of-function lines to ensure that the differences we detect would not be solely due to tissue-culture induced changes in methylation.

**Figure 2.**
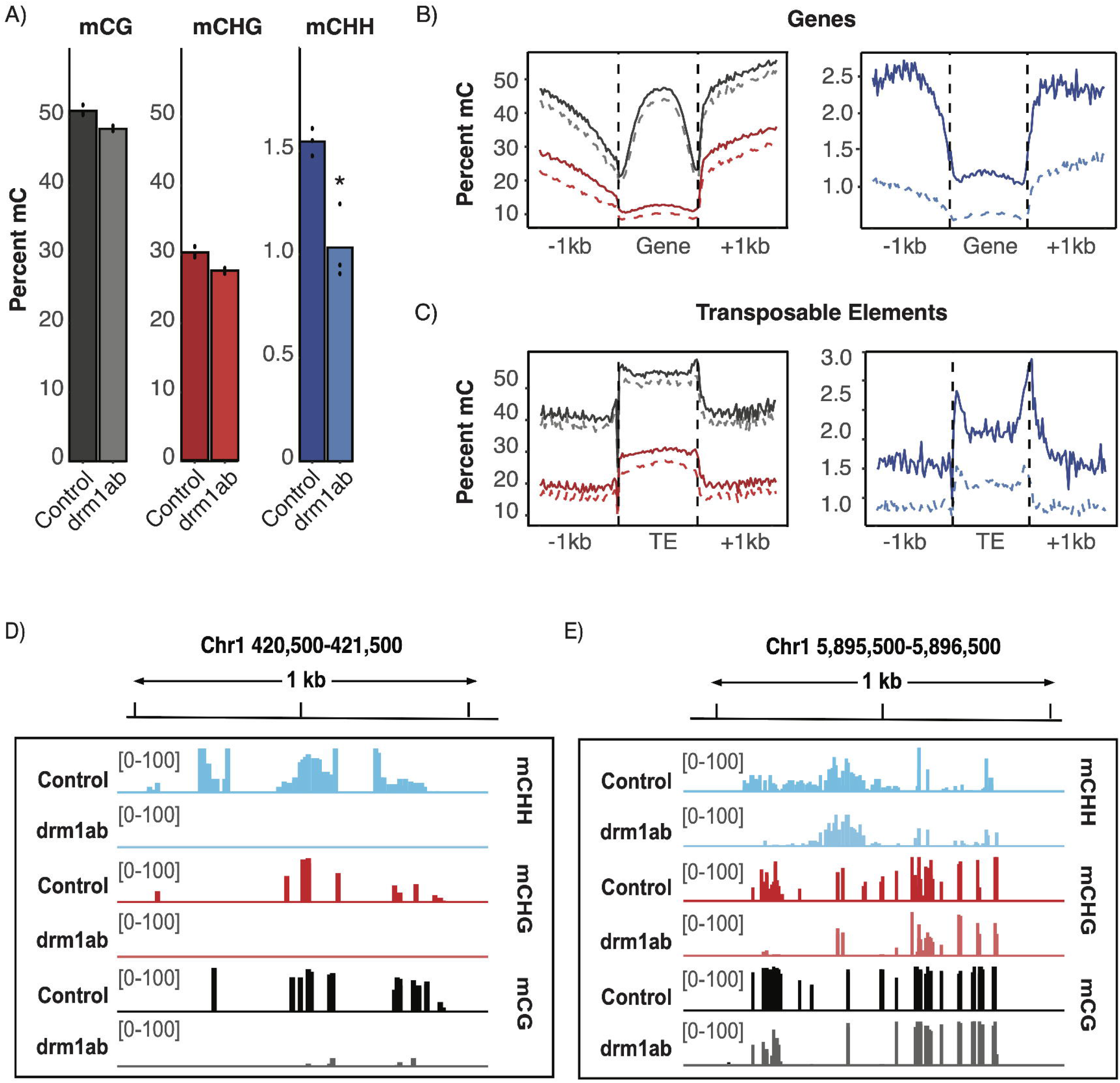
DNA methylation changes in *drm1ab* plants. (A) Genome-wide mean mCG, mCHG and mCHH levels were assessed in three biological replicates of nonedited tissue-culture control plants and *drm1ab* plants. Asterisk indicates significantly lower levels of mCHH methylation in *drm1ab*. (B) Metaplots of mCG, mCHG or mCHH levels in genic regions. Solid lines show the profile in tissue-culture control plants, dashed lines show the levels in *drm1ab*. A different scale is used for mCHH due to overall lower methylation levels in this context. Genes are all oriented 5’ to 3’ and the dashed lines indicate the genic region normalized to the same length. The region to the left or right of the dashed vertical lines include 1kb of upstream or downstream sequences. (C) Similar metaprofiles of mC levels within and surrounding structurally annotated transposable elements. Genome viewer snapshots showing an example of (D) DRM-dependent mC loss and (E) DRM-independent maintenance of mC.

### Widespread loss of CHH methylation in drm1ab mutant plants

Context-specific DNA methylation levels were evaluated genome-wide (Figure 2A) and using metaprofiles over genes or TEs (Figure 2B-C). This revealed substantial loss of CHH methylation in *drm1ab* relative to the control. CHH methylation is lost in regions that exhibit elevated levels of methylation including the regions surrounding genes as well as within TEs (Figure 2B-C). However, there is still CHH methylation remaining in the *drm1ab* plants, especially within TEs. A visualization of several genomic regions revealed that regions of high CHH methylation in the tissue culture control plants can be divided into regions that require DRM (DRM-dependent) and regions that have CHH methylation that is not dependent upon DRM (DRM-independent) (Figure 2D). The proportion of CHH methylation that is lost in *drm1ab* was assessed for genomic regions with varying levels of CHH methylation in the control (Figure S2C). The vast majority of regions with high (>20%) CHH have significant losses of methylation in *drm1ab* plants (Figure S3C). In regions with moderate levels of CHH methylation (5%-20%) we find that some of these have strong losses of methylation and other regions do not lose methylation. Interestingly, regions with low, but detectable levels (2-5%) of CHH methylation rarely lose methylation in the *drm1ab* plants and these regions are more prevalent in the genome than regions with high (>20%) CHH methylation (Figure S3C). The loss of DRM activity results in strong loss of CHH methylation at regions with high CHH methylation but rarely affects the large number of regions with low levels of CHH methylation.

To gain a better understanding of CHH methylation changes, and how these are related to CG and CHG methylation, we identified all 100bp tiles with >20% CHH methylation in the control sample. The methylation levels of these tiles were assessed in *drm1ab* to identify DRM-dependent tiles (>80% methylation loss in *drm1ab*), DRM-intermediate tiles (20-80% methylation loss in *drm1ab*), and DRM-independent CHH tiles (<20% methylation loss in *drm1ab*) (Table 1). The CG, CHG and CHH methylation levels in both control and *drm1ab* were evaluated at these regions (Figure 3A-B). The majority (79%) of regions with CHH levels >20% in wild-type are DRM-dependent with only 4% that are DRM-independent (Table 1). We observed that DRM-dependent CHH methylation is often accompanied by high levels of CG and CHG methylation (Figure 3A). In *drm1ab*, the CHG methylation is lost in the vast majority of these regions and the CG methylation is reduced at some loci, but not at others (Figure 3A). The regions of the genome where we observed DRM-independent CHH methylation generally have high levels of CG and CHG methylation that are not dependent upon DRM (Figure 3B).

**Table 1.**
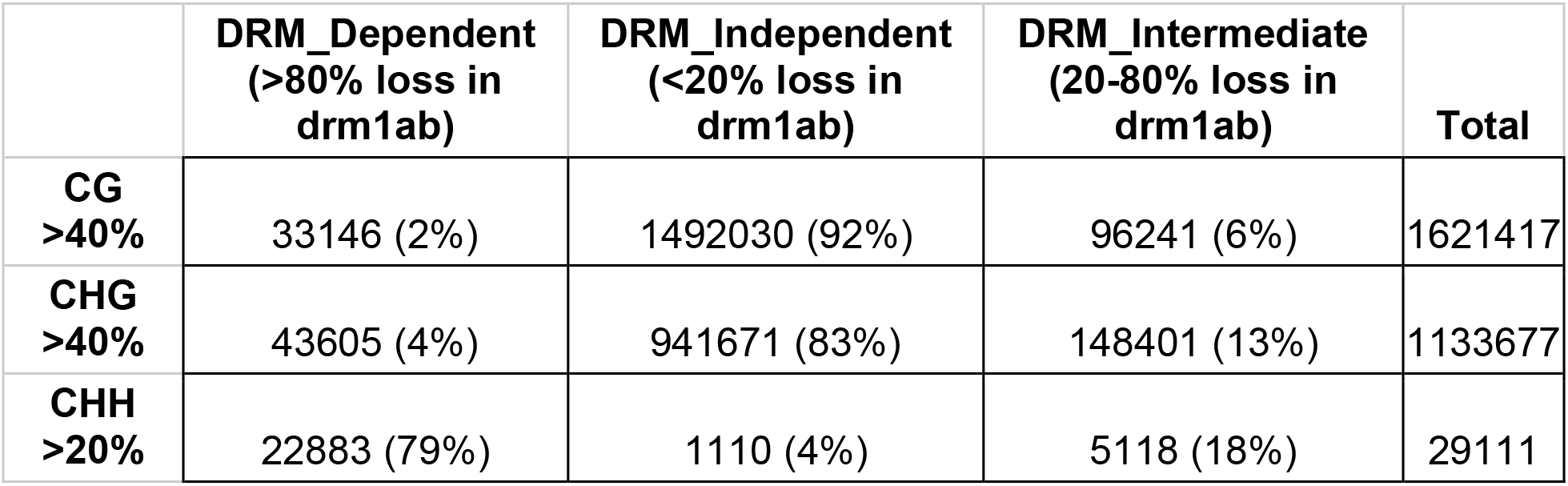
Count of DRM-dependent, DRM-intermediate, and DRM-independent methylated tiles in all contexts.

|  | **DRM_Dependent**<br>**(>80% loss in**<br>**drm1ab)** | **DRM_Independent**<br>**(<20% loss in**<br>**drm1ab)** | **DRM_Intermediate**<br>**(20-80% loss in**<br>**drm1ab)** | **Total** |
| --- | --- | --- | --- | --- |
| **CG**<br>**>40%** | 33146 (2%) | 1492030 (92%) | 96241 (6%) | 1621417 |
| **CHG**<br>**>40%** | 43605 (4%) | 941671 (83%) | 148401 (13%) | 1133677 |
| **CHH**<br>**>20%** | 22883 (79%) | 1110 (4%) | 5118 (18%) | 29111 |

**Figure 3.**
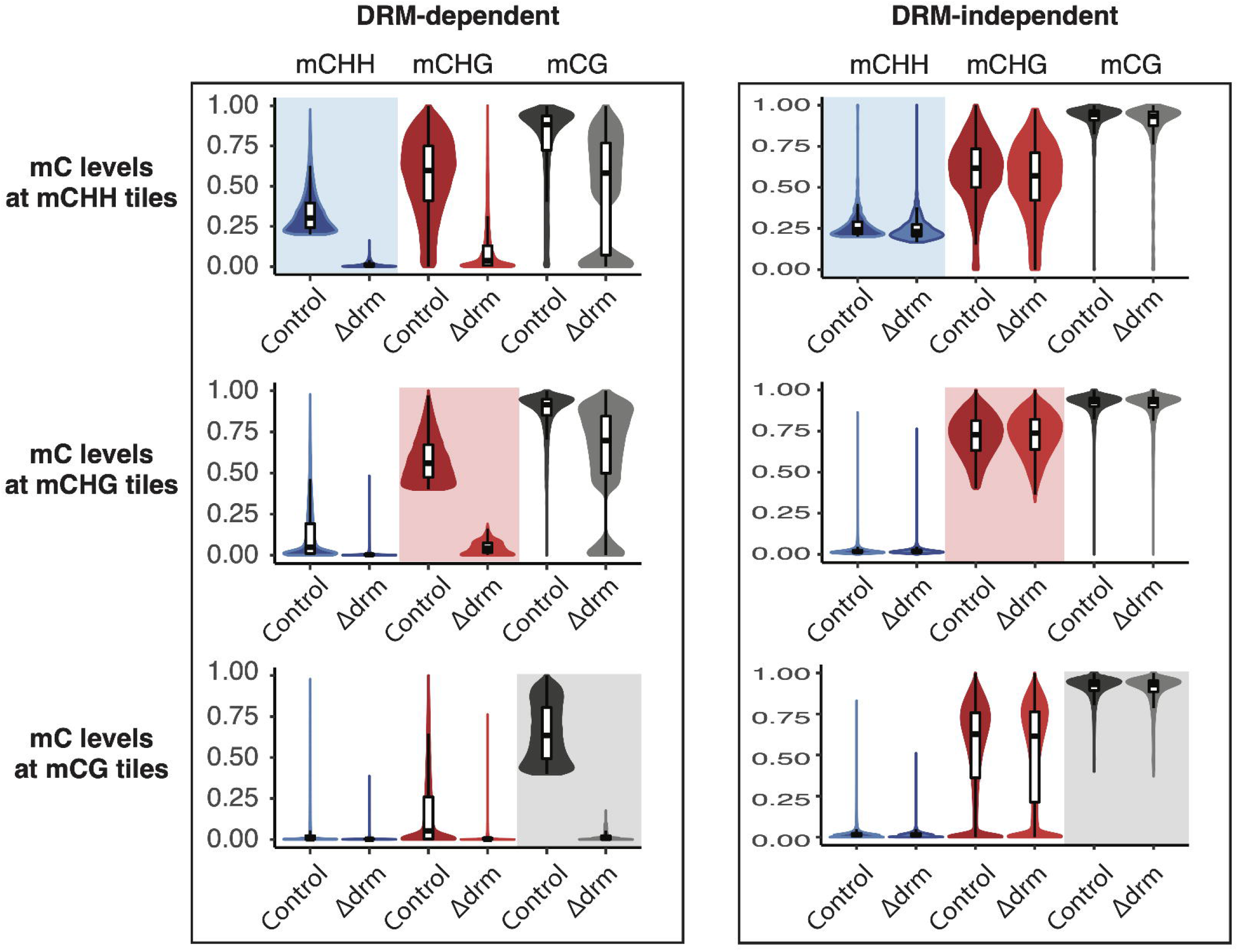
Comparisons of context-specific mC levels at DRM-dependent and DRM-independent loci. 100 bp tiles classified as methylated (>= 40% for CG or CHG, >= 20% for CHH) were examined to see if methylation was lost in *drm1ab* plants. DRM-dependent tiles lost >80% mC while DRM-independent tiles lost <20% mC in *drm1ab*. Each violin plot includes context-specific DNA methylation levels in both mutant and control plants for the set of DRM-dependent or -independent tiles. Shaded boxes highlight the mC context used to select the subset of tiles included in each plot. Details of tile classification can be found in table 1.

DRM is expected to function in the RdDM pathway and to primarily contribute to maintenance of CHH methylation. However, genome-wide levels of CG and CHG methylation show 2-6% reductions (Figure 2A-B). Methylation levels in all contexts were determined at each 100 bp genomic tile in the control and *drm1ab* samples. Differentially methylated tiles were classified as either hypermethylated (higher methylation in *drm1ab*) or hypomethylated (lower methylation in *drm1ab*) (Figure S4). In all three contexts there are many more examples of hypomethylated tiles *drm1ab* relative to the control (Figure S4). All genomic regions with high (>40%) CG or CHG methylation levels in the control sample were evaluated to determine what proportion are DRM-dependent (Table 1). In contrast to high CHH regions in which the majority are dependent on DRM, only a small proportion of these high CG or CHG methylated regions are affected in *drm1ab*. However, since the number of genomic regions with high CG or CHG methylation vastly outnumbers the regions with elevated CHH, there are more total CG or CHG hypomethylated tiles genome-wide (Table 1).

The *drm1ab* mutant might be expected to affect CG and CHG methylation in regions with high CHH methylation, but we did not expect substantial changes in CG and CHG methylation at regions without CHH methylation. We sought to determine whether the changes in CG and CHG methylation in *drm1ab* co-occurred with changes in CHH methylation or whether some CG and CHG methylation losses occurred in regions without CHH methylation. We found many examples of CG or CHG DRM-dependent hypomethylation at tiles with low or no CHH methylation (Figure 3C-F). A comparison of the regions with CG and CHG methylation loss found some examples of dual loss in both contexts as well as many with specific loss in CG or CHG (Figure 4A). The CG, CHG or CG/CHG methylation losses were then evaluated to assess what proportion overlap, are within 300bp of, or are greater than 300bp from a CHH DRM-dependent or CHH DRM-intermediate tile (Figure 4A and B). A portion of the DRM-dependent CG or CHG methylation is found in regions that have moderate or high levels of CHH that are DRM-dependent or DRM-intermediate (Figure 4A-B). In addition, another 7-10% of the CG or CHG dependent methylation occurs within 300 bp of a region with CHH methylation loss in *drm1ab* (Figure 4B). This suggests that the loss of a small region of CHH methylation can result in broader loss of CG and/or CHG methylation at some loci (example in Figure S5). The mechanisms leading to losses of CG or CHG methylation in *drm1ab* in regions distal to CHH methylated regions are unclear.

**Figure 4.**
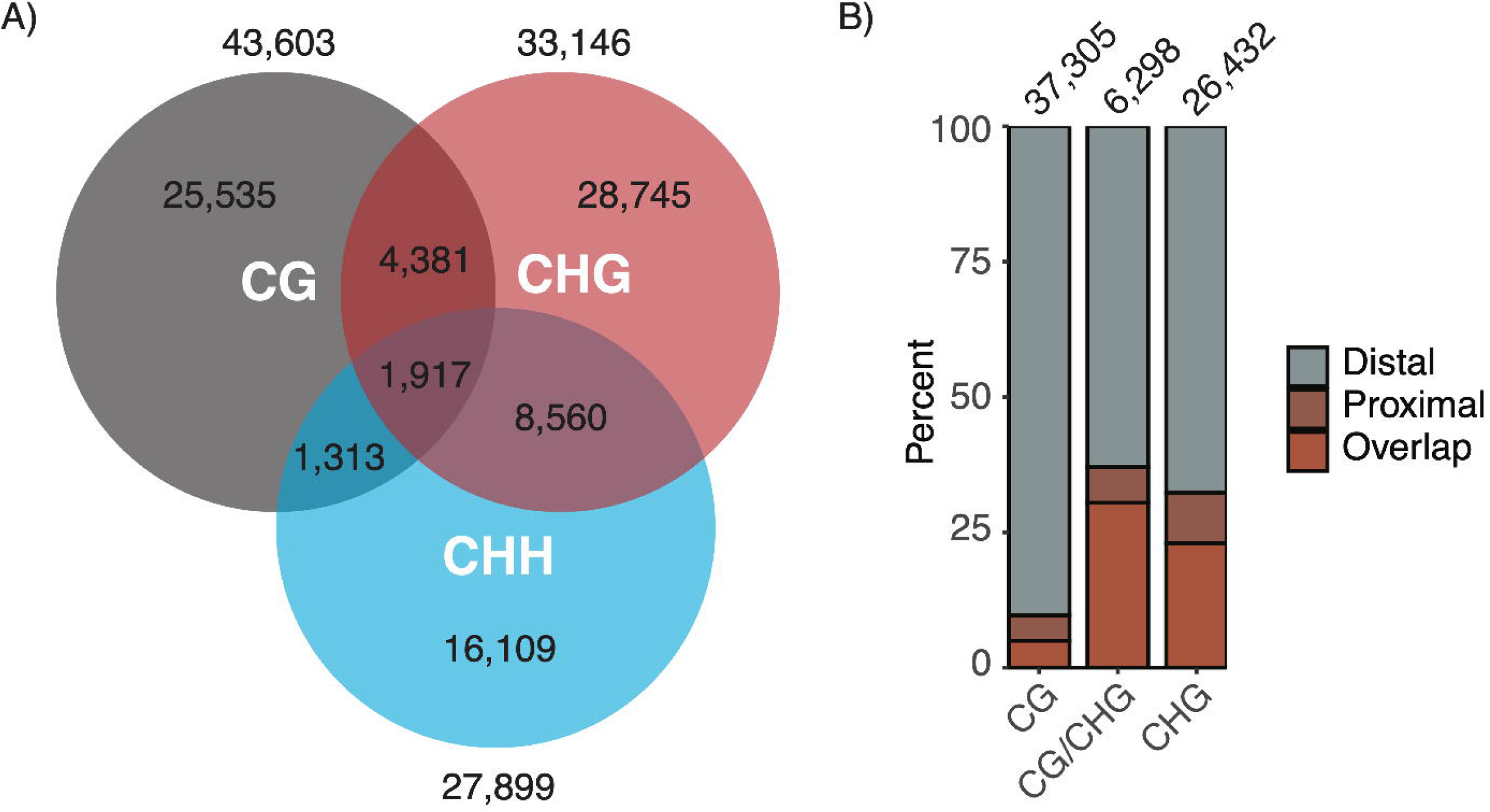
Relationship of DRM-dependent mCG and mCHG losses to mCHH hypomethylated tiles. (A) Venn diagram of the overlap of tiles with mCG and/or mCHG DRM-dependent methylation with tiles with mCHH DRM-dependent or -intermediate loss. (B) The proportion of mCG, mCHG, or mCG/CHG DRM-dependent tiles that overlap, are proximal to (within 300 bp), or distal to (greater than 300 bp) mCHH hypomethylated tiles. The number of hypomethylated tiles in each mC context can be found in Figure S3.

### Limited changes in gene expression in DRM mutants

Transcriptome profiling was performed using RNAseq on the same seedling tissue samples used for WGBS with the addition of multiple replicates for wild-type ME034V. Principal component analysis (PCA) showed limited variation between wild-type ME034V and tissue culture derived ME034V plants, and, as expected based on the PCA result, there are very few differentially expressed genes between these samples (Figure S6, 5A). However, PCA clustering and differential gene expression analysis finds evidence for hundreds of gene expression changes in *drm1ab* plants (Figure 5A-B). The *drm1ab* plants have more genes that are up-regulated compared to the control samples, including many genes with >10-fold up-regulation (Figure 5A-B). The observed changes in expression in *drm1ab* plants may represent direct effects of changes in DNA methylation on gene expression or could represent secondary effects due to a small number of direct targets that influence expression of other genes. In order to identify potential direct effects of loss of RdDM we initially focused on the subset of genes with CHH methylation immediately over the TSS region. We identified 1,043 genes with CHH methylation (>20%) in the region immediately surrounding the annotated transcription start site (TSS) in tissue culture control plants. Many (529) of these genes that contain high CHH immediately at or surrounding the TSS are expressed in the control, suggesting that the presence of CHH at or near the TSS is not necessarily silencing gene expression. While the vast majority of these genes (98%) are hypomethylated in *drm1ab*, only 3.5% are differentially expressed (24 up in *drm1ab*, 12 down in *drm1ab*) (Figure 5C). Over 95% of the genes with elevated CHH methylation surrounding the TSS do not exhibit changes in expression in *drm1ab*. These observations suggest that there are relatively few genes that are direct targets for silencing by RdDM in seedling leaf tissue of *Setaria viridis*.

**Figure 5.**
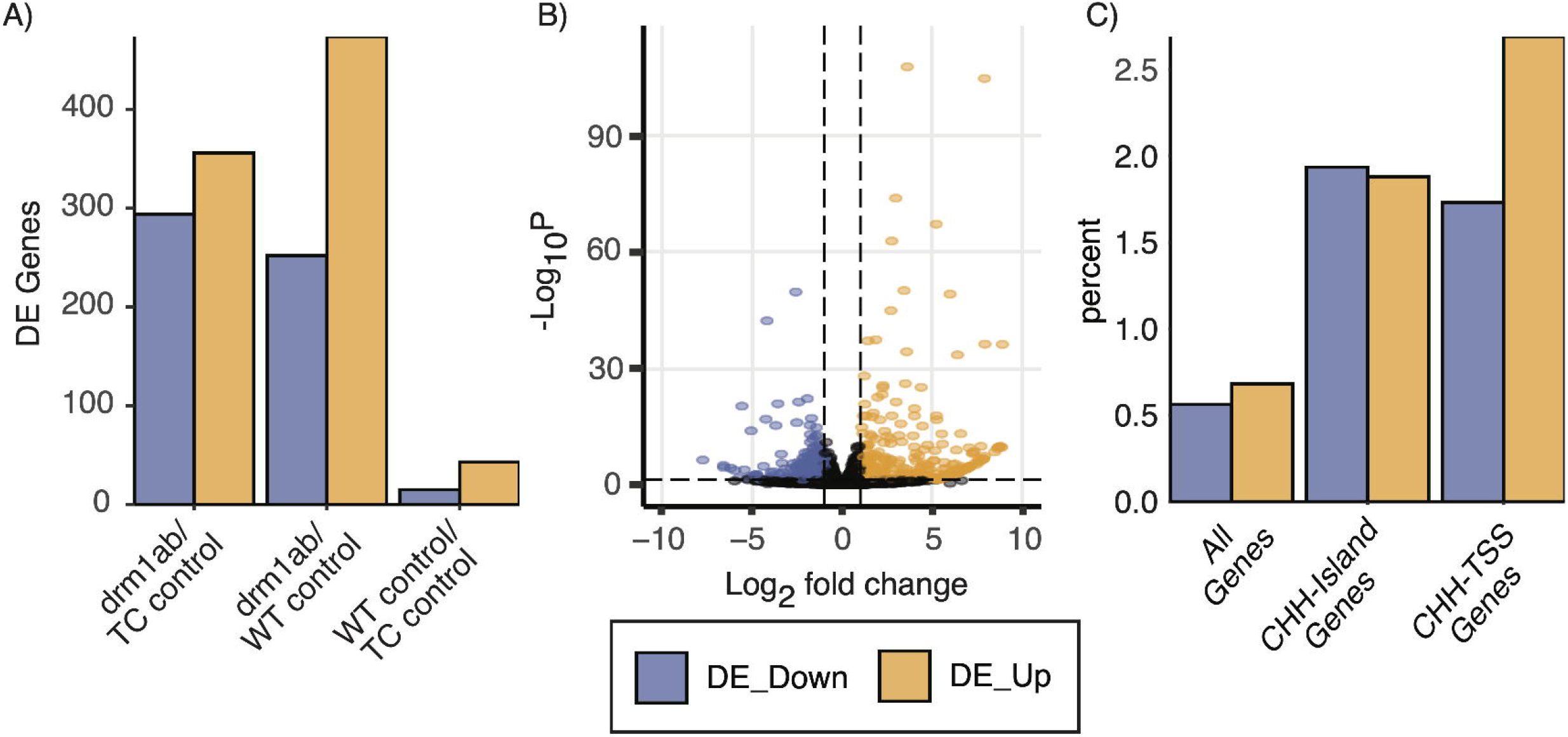
Transcriptome changes in *drm1ab* plants. (A) The number of genes with significant differences in expression (padj<0.05; >2x fold-change) was determined for all contrasts. (B) A volcano plot showing magnitude and padj for differentially expressed genes in the drm1ab:tissue culture comparison. Significant differences are indicated using blue and orange data points. (C) The proportion of genes that are up-(orange) or down-regulated (blue) is shown for all genes, the sub-set of 5,424 genes that contain a tile of >20% CHH within 1kb of the TSS (CHH-Island genes), or the 1,043 gene with >20% CHH methylation in the 100bp tile directly over the TSS or the two adjacent tiles (CHH-TSS genes).

Previous analysis of DNA methylation in monocots has shown that methylated CHH regions (mCHH islands) are often found upstream of highly expressed genes (Niederhuth *et al*., 2016; Li *et al*., 2015; Gent *et al*., 2013). It is unclear whether the mCHH islands influence nearby gene expression or if open chromatin associated with gene expression enables DRM-dependent methylation of these regions. We sought to determine if losses of CHH methylation at mCHH islands in *drm1ab* resulted in changes in expression of these genes. We classified 5,424 genes as having an mCHH island (requires at least one 100bp tile with >20% CHH methylation in the 1kb promoter region). Many (3,171) of these genes are expressed in control conditions and these genes tend to be higher expressed than genes without mCHH islands as previously observed in maize (Li *et al*., 2015; Gent *et al*., 2013). The genes containing mCHH islands exhibit only slightly higher proportions of DEGs than in all genes and there are similar proportions of up- and down-regulated genes in *drm1ab* relative to control (Figure 5C). Over 95% of the genes with an mCHH island do not show altered expression in the *drm1ab* plants, suggesting that the presence of mCHH islands in gene promoters has limited functional significance for gene expression levels in seedling leaf tissue.

### Identification of transposable elements that are up-regulated in drm1ab mutants

RdDM has been shown to play important roles in maintaining silencing of transposable elements in Arabidopsis (Cao *et al*., 2003; Tran *et al*., 2005; Stroud, Greenberg, *et al*., 2013; Chan *et al*., 2004; Cao and Jacobsen, 2002a). However, in Arabidopsis and other plants it has been shown that there is often redundant control of transposable element (TE) silencing through multiple DNA methylation pathways and sole loss of CHH methylation only results in limited TE activation (Chan *et al*., 2006; Henderson and Jacobsen, 2008; Stroud *et al*., 2014). We sought to investigate whether there are *Setaria* TEs that are transcriptionally activated in the *drm1ab* plants. The Extensive de-novo TE annotator (EDTA) (Ou et al. 2019) pipeline was used to perform both a structural and a homology based annotation of TEs in the *S. viridis* ME034V genome. Two distinct approaches were used to monitor expression of TEs. The first approach assessed uniquely aligned RNAseq reads that align to regions annotated as TEs using the structural annotation of intact ME034V TEs. There were 454 TEs with detectable expression (>5 uniquely mapping reads) in either tissue culture or *drm1ab* samples and 33 of these were differentially expressed (padj<.05 and greater than 2-fold change) between tissue culture and *drm1ab* samples (Figure 6A). The majority (25/33) of the differentially expressed TEs were up-regulated in *drm1ab* and 14 of these had little or no expression in the control plants indicating the requirement for RdDM to maintain effective silencing of these TEs. The up-regulated TEs include 11 class I retrotransposons as well as 14 class 2 terminal inverted repeat and helitron DNA transposons (Table S1). This approach was useful to identify TEs that are up-regulated, but has two significant limitations. First, the reliance on uniquely mapping reads limits detection of TE families with multiple highly similar elements which might be common in TE families regulated by RdDM. Second, in many cases the expression that was detected for TEs likely reflects partial transcripts rather than expression of the full-length TE.

**Figure 6.**
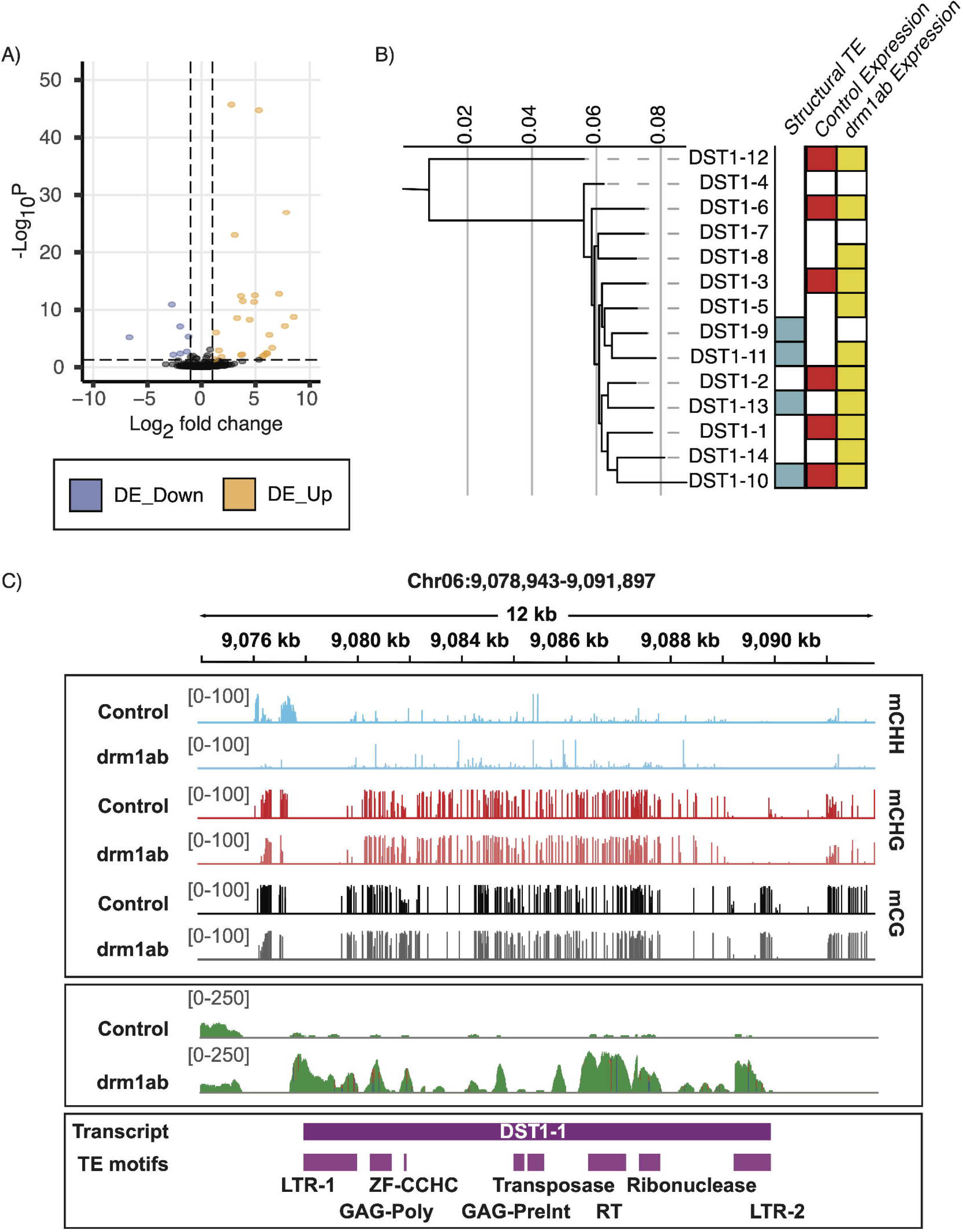
Expression of structurally intact TEs and analysis of DST-1-like TEs. (A) A volcano plot showing magnitude and padj for differentially expressed structurally annotated TEs in the tissue culture control:drm1ab comparison. Significant differences are indicated using blue and orange data points. (B) A maximum likelihood relatedness tree generated from an alignment of the putative transcripts of the 14 DST1-like TEs. DSTs that: overlap a structurally annotated TE, have expression in control, and/or drm1ab are indicated. (C) Genome browser snapshot of the DST1-1 region showing mC levels in tissue-culture and drm1ab plants, (D) RNAseq coverage from tissue-culture control and drm1ab plants, and (E) the location of the predicted DST1-1 transcript and TE-associated domains identified by CD-BLAST.

The second approach to monitor TE expression was implemented using a *de novo* transcriptome assembly of the RNAseq reads from the wild-type and *drm1ab* plants. This enables the identification of TEs that generate potential full-length transcripts. *De novo* transcriptome assembly does not rely upon alignment to the reference genome and therefore can potentially identify transcripts that arise from repetitive sequences. The RNAseq reads from the wild-type, tissue culture wild-type, and *drm1ab* samples were aligned to *de novo* assembled transcripts that were greater than 1kb in length to identify 103 up-regulated transcripts in *drm1ab* (minimum 2-fold change and padj<0.05). These transcripts include both gene and TE sequences. To focus on putative TEs, we removed any of the 103 *drm1ab* up-regulated transcripts with >50% overlap of an annotated gene based on alignment of the transcripts to the genome. There were 29 up-regulated transcripts that do not align to annotated genes. The analysis of conserved domains within the putative ORFs of these transcripts identified five of these transcripts that contain TE-associated domains and three additional transcripts that overlap structurally annotated TEs. We refer to these eight transcripts as DRM Silenced TEs (DSTs) (Table S2). The eight DSTs were aligned to the genome to identify the best matching genomic sequence and we assessed the presence of LTR/TIRs and target site duplications. Only DST1 contained intact structural features that would be necessary for active transposition and we focused on further characterization of this element and related family members.

The DST1 *de novo* assembled transcript is 8.7kb in length and is >8 fold up-regulated in *drm1ab* (Table S2). Alignment of the *de novo* assembled DST1 transcript to the genome assembly of ME034V revealed 15 highly similar sequences (Table S3). Four of the DST1 elements are identified as TEs in the structural annotation and another 10 are at least partially annotated as TEs in the homology-based annotation. These annotated TEs were not identified as differentially expressed based on alignments of RNAseq reads to the genome, likely due to the repetitive nature of the family and limited uniquely mapping reads. The internal (non-LTR) sequences typically have 96-98% identity between members of this family with one element (DST1-12) having lower similarity, indicating that this may be the oldest member of the family. A phylogeny of the DST1 family based on the internal alignable sequences does not reveal any subgroups with very close relationships that would indicate on-going movement of this set of elements (Figure 6B). The majority of these (14/15) have intact LTR sequences on the 5’ and 3’ end with 90-96% identity of the two LTRs suggesting that these elements do not represent particularly recent transposition events. However, there is substantial variability in the LTR sequences between different elements of the family. Only DST1-2/DST1-11 and DST1-4/DST1-6 share similar LTRs. A comparison among all other elements revealed that there is conservation (90-95%) identity at the first 250-400bp and in the last 1.2kb of the ~2kb LTRs. The middle region of 300-800bp is highly variable and can not be aligned among family members. The DST1-1 element only contains homology for the first 250bp region. DST1 is an intriguing LTR family with highly conserved internal sequences but a variable region within the LTR reminiscent of observations for the Tnt1 TE family in tobacco (Casacuberta et al. 1997).

Alignment of RNAseq reads to the DST1 transcript reveals an ~8-fold increase in expression in the *drm1ab* individuals. However, many of the aligned reads contain SNPs relative to the DST1 assembled transcript, potentially reflecting expression of multiple family members with slight sequence variation. The transcripts that arise from DST1 elements were assessed to determine if the activation in *drm1ab* occurs at a single element or reflects coordinate activation of several members of the family. Based on SNPs within the DST1 sequence there is evidence for the expression of 11 members of the DST1 family and five of the DST1 elements are only detected in the *drm1ab* mutant (Figure 6B). The DST1 family members are often highly methylated with high CHH levels (>75%) at, or near, the LTRs and six of the DST1 elements have loss of CHH methylation at or within 1 kb of the annotated element in *drm1ab* (Figure 6C).

## Discussion

The DRM methyltransferases are critical for RdDM in plants. However, there are variable consequences for the loss of functional RdDM among different plant species. While there are some gene and TE expression differences in *Arabidopsis thaliana*, there are limited phenotypic differences (Cao and Jacobsen, 2002a; Chan *et al*., 2006). In contrast, rice and maize mutants lacking functional RdDM exhibit developmental abnormalities (Moritoh *et al*., 2012; Sidorenko *et al*., 2009; Alleman *et al*., 2006). In this study we isolated *S. viridis* plants with loss-of-function mutations in both catalytically active *DRM* genes. While there are some phenotypic differences such as reduced stature, the plants are fully viable with normal inflorescences. This suggests that a functional RdDM pathway is dispensable under standard growth conditions. However, it is worth noting that we have only maintained the *drm1ab* mutants in the homozygous mutant condition for several generations. One important function of RdDM might be to ensure the faithful maintenance and inheritance of DNA methylation patterns. Loss of fidelity in maintaining heterochromatin may have growing phenotypic consequences after many generations in the absence of RdDM activity.

A detailed analysis of the seedling leaf methylome in *drm1ab* plants revealed significant changes in CHH methylation, as expected. In particular, we find loss of CHH methylation at the vast majority of genomic loci with high (>20%) CHH methylation in wild-type plants. In contrast, many genomic regions that had lower, but detectable, levels of CHH methylation (5-10%) are not changed in a *drm1ab* mutant. It seems that the RdDM pathway is responsible for high levels of CHH methylation that is often found at the edges of TEs, especially near genes. In contrast, the lower levels of CHH methylation are often found within larger TEs and this methylation is likely the result of *CMT2* or other chromomethylases similar to observations in other plant species (Zemach et al. 2013).

The analysis of CG and CHG methylation patterns in *drm1ab* plants revealed some unexpected findings. First, we found that CHG methylation was rarely maintained at loci that had lost CHH methylation. The regions with high CHH methylation typically had CHG methylation in wild-type plants and both CHH and CHG methylation were lost in the *drm1ab* mutant. This suggests widespread failure to maintain CHG methylation at RdDM targets in the absence of functional DRM. Second, we found numerous examples of CG and/or CHG methylation loss in regions that were not located at or near loci with high CHH methylation. It is not clear why functional DRM is necessary for maintenance of CG/CHG methylation at these loci. One possible explanation is that these regions have elevated CHH at other developmental stages and the loss of active RdDM at that stage results in the loss of CG/CHG methylation which is observed in leaf tissue. It is also possible that these sites are targets of active DNA demethylation and require RdDM for maintenance.

There are relatively few changes in gene expression in *drm1ab* plants. We hypothesized that the subset of genes with high CHH methylation over the TSS would be up-regulated in *drm1ab*. However, we found that many of these genes are already expressed in wild-type plants that contain methylated TSS regions and very few of these genes have altered expression when the methylation is lost. This observation suggests a complex relationship between RdDM activity and gene expression. It is possible that many of these genes are redundantly silenced by RdDM activity as well as CMT2/3 and *MET1* maintenance activities. It is possible that loss of CHH methylation may destabilize silencing but does not lead to activation.

A small set of TEs that are silenced in wild-type plants exhibit transcriptional activity in *drm1ab* plants. These TEs appear to require RdDM for full silencing. We identified a novel TE family that exhibits coordinated activation of multiple loci in *drm1ab* mutants. This family will be of particular interest in future studies of potentially active TEs in the Setaria genome.

## METHODS

### Plant material and growth conditions

*Setaria viridis* variety ME034V was used in this study. The tissue culture wild-type control plants and the *drm1ab* plants were derived from the same T0 transgenic plant as described in the previous study (Weiss et al. 2020). Dormancy was broken by incubating freshly harvested seeds at 29°C for 24 h in a 1.4 mM gibberellic acid and 30 mM potassium nitrate solution (Sebastian et al. 2014). Seeds were then sterilized with 50% bleach for 10 min, rinsed five times with water, and then planted on germination media [0.5X MS, 0.5% sucrose, 0.4% Phytagel (Sigma, St Louis, USA), pH 5.7]. 6 days after germination, seedlings were transplanted to soil and grown under a 16:8 h light/dark photocycle at 26°C/22°C (day/night) and 30% relative humidity, according to a modified protocol from Huang et al.

### Guide RNA design and vector construction

The genomic sequences of *DRM1a* and *DRM1b* were obtained prior to the publication of the ME034V genome and were identified by BLAST searching the *S. viridis* A10.1 reference (Mamidi et al. 2020) using the phytozome database (https://phytozome.jgi.doe.gov). CRISPR gRNAs were designed to target the conserved domains in each gene using CRISPOR (Haeussler et al. 2016). Conserved domains were identified by aligning the coding sequences from *S.viridis* with orthologs from brachypodium, maize and Arabidopsis. Construction of the T-DNA construct, pTW045, was described previously using the Golden Gate assembly method (Weiss et al. 2020)

### T-DNA transformation and tissue culture

Agrobacterium tumefaciens-mediated transformation of *S. viridis* ME034Vwas performed as described previously with a few modifications (Van Eck et al. 2017; Weiss et al. 2020). Callus initiation was first performed by removing the seed coats and sterilizing seeds with a 10% bleach plus 0.1% Tween solution for 5–10 minutes under gentle agitation. Seeds were then placed on callus induction media with the embryos facing upward at 24°C in the light for 1 week and then moved to dark for callus initiation. Embryogenic calli were collected after 4–7 weeks and inoculated with the AGL1 strain harboring the T-DNA construct pTW045 (Weiss *et al*., 2020). Inoculated calli were placed on a co-culture medium and incubated in the dark at 20°C for 5–7 days. Transformed calli were transferred to selection medium with 50 mg L^-1^ hygromycin for 4 weeks at 24°C. Selected calli were subcultured on plant regeneration media with 20 mg L–1 hygromycin with 16-h light to allow the growth of transformed shoots. Elongated shoots were transferred to rooting medium with 20 mg L^-1^ hygromycin. Shoots were transplanted to soil and grown to maturity.

### Genotyping and *drm1ab* identification

The *drm1ab* and wild-type tissue culture control plants were identified using genomic PCR with restriction enzyme digestion (CAPS assay) followed by Sanger sequencing. PCR was performed with GoTaq Green Master Mix (Promega Corp., Madison, WI, USA) in accordance with the manufacturer’s instructions, with an annealing temperature of 58°C and an extension time of 1 min. Amplicons were then subjected to restriction enzyme digestion using an enzyme that overlaps with the CRISPR-Cas9 cleavage site. PCR amplicons made with the corresponding primers were subjected to Sanger sequencing. T-DNA transgene detection was conducted using two methods: genomic PCR amplification of the hygromycin gene that is close to the T-DNA left border and a luciferase assay to detect the expression of the luciferase reporter gene that is next to the T-DNA right border. The luciferase assay procedure was conducted using the Bio-GloTM Luciferase Assay System (Promega Corp.) in accordance with the manufacturer’s instructions. All the primer sequences used in the present study can be found in Table S4.

### Methylome profiling

DNA was extracted from tissue collected from the 3rd and 4th leaf of 2.5 week old *S. viridis* plants grown in a growth chamber with 31°C/21°C 12-hour/12-hour day/night conditions. For each sample, tissue from 3-4 plants were pooled prior to CTAB DNA extraction. In total, seven samples from pooled tissue were converted and analyzed: one, three biological replicates of unedited plants regenerated from tissue culture, and three biological replicates of *drm1ab* edited plants. The samples were converted for sequencing with the NEBNext enzymatic methyl-seq kit (NEB) and sequenced at the University of Minnesota Genomics Center. All samples were multiplexed in a full Novaseq S1 lane with 150bp paired-end sequencing. Sequencing reads were trimmed with Trim galore! version 0.4.3, powered by cutadapt v1.8.1 (Martin 2011) and fastqc v0.11.5 and aligned to the ME034V reference genome (Thielen *et al*., 2020) with bsmap v2.74 using the following parameters: -v 5 -r 0 -p 8 -q 20 (Xi and Li, 2009).

The genome was divided into adjoining 100-bp tiles and each tile was classified as one of: “missing data” (including “no data” and “no sites”), “CHH > 15%”, “CG/CHG”, “CG-only”, “unmethylated”, or “intermediate” following the classifiers and hierarchy outlined in (Crisp *et al*., n.d.). The ratio of tiles in each category is displayed in Figure S2. We further classified tiles as DRM-dependent, DRM-independent, or DRM-intermediate. Tiles classified as “missing data” in either the tissue culture or *drm1ab* mutant samples were omitted from this analysis. For all remaining tiles, we first asked whether the tile was methylated in one or more contexts in the tissue-culture control plant. If yes, the percentage of methylation loss in *drm1ab* was determined. Loss of >80% methylation was classified as DRM-dependent, loss of 20-80% was classified as DRM-intermediate, and a change of <20% as DRM-independent (Table 1).

In order to determine the relationship of CHH methylation with gene expression, all genes were classified as either CHH-TSS genes and/or CHH-Island genes. CHH-TSS genes have a tile with >20% mCHH in the tissue culture samples that is overlapping, or within one tile, of the annotated transcriptional start site. CHH-Island genes have at least one tile with >20% mCHH in the tissue culture sample that is between 100 and 1000-bp upstream of the TSS. It is possible for a gene to be both a CHH-TSS gene and a CHH-Island gene. In addition to association with gene expression, we were interested in whether DRM-dependent CG and/or CHG methylated tiles were often found at or near CHH DRM-dependent and CHH DRM-intermediate tiles. For this, we classified each CG, CHG, and CG/CHG DRM-dependent hypomethylated tile as overlapping, within 300-bp (proximal), or greater than 300-bp (distal) from a CHH DRM-dependent/intermediate tile (Fig 4).

### Transcriptome profiling

Prior to DNA extraction, a portion of the ground tissue used for methylome profiling was saved for RNA extraction with the QIAGEN RNeasy mini kit (QIAGEN). For RNAseq, two additional wild-type biological replicates were included. RNA was submitted to the UMGC facility for 150 bp cDNA paired-end library preparation and run on the Illumina NovaSeq 6000. Sequencing reads were trimmed as described in the above methylation profiling section. The ME034V genome was indexed using the –runMode geneomeGenerate command of STAR 2.7.1 (Dobin et al., 2013) using an annotation file that included the primary transcript for each gene as well as all structural TEs. The trimmed reads were aligned to the indexed genome with the –quantMode GeneCounts feature of STAR 2.7.1 (Dobin *et al*., 2013).

Read counts were imported into R v4.1.1 (R Core Team, 2021). Normalization (median of ratios) and differential expression was determined using DEseq2 (Love *et al*., 2014). A gene or structural TE was determined to be differentially expressed if the absolute value of the log2 fold change was greater than 1 and the adjusted P-value was less than 0.05. EnhancedVolcano (https://github.com/kevinblighe/EnhancedVolcano) was used to visualize differentially expressed genes and TEs.

Additional RNAseq datasets were used to compare our observed expression ratios of *Drm1a* to *Drm1b* and *Drm3* with previously published data (Fig S1). The data for *S. viridis* cultivar A10 were downloaded from the Phytozomev13 portal (Goodstein *et al*., 2012). The data for ME034v are from (Thielen *et al*., 2020).

### De novo Transcript assembly and analysis

A *de novo* transcriptome assembly was generated using pooled trimmed RNAseq reads from all nine samples with Trinity version 2.10.0 (Haas *et al*., 2013). The minimum contig length was set to 200 bp. The *de novo* transcripts were indexed with the ‘gmap_build’ command of gmap version 2015-09-26 (Wu and Watanabe, 2005). Default ‘gmap’ parameters were used to map the *de novo* transcripts back to the ME034V reference genome.

Next, the RNAseq reads from each sample were mapped to the transcriptome assembly with Salmon version 1.2.1 (Patro *et al*., 2017) using default parameters in order to determine transcripts per million (TPM) in individual biological samples. Differential transcript expression was determined as previously described using the Salmon TPM data as input.

To find transcripts from the transcriptome assembly that may represent TEs that are upregulated in the *drm1ab* mutant, we first filtered our DE transcript list to only include transcripts greater than 1-kb, reasoning that any shorter transcripts would not encode functional TEs. Next, we set a DE threshold of 2-fold upregulation in the mutant with an adjusted P-value < 0.05. Transcripts with at least 50% of their length annotated as genes based on overlap of coordinates from the gmap alignment and the current ME034V gene annotation were removed. The remaining transcripts were subjected to conserved domain BLAST (Lu *et al*., 2020). Transcripts without TE-associated domains (e-value < 0.01) were omitted from further analyses. Finally, the transcript plus the 3-kb flanking sequence on either side was submitted for a self vs. self BLAST to determine if putative LTR or TIR sequences could be observed.

Of the five transcripts that had evidence for upregulation, one had detectable LTR sequences (now referred to as DST1). DST1 family members were identified in the ME034V genome using BLAST (Altschul et al., 1990)with parameters ‘blastn -perc_identity 75 -qcov_hsp_perc 75’. A total of 15 similar sequences were identified and classified as a TE family using the 80-80-80 rule (Seberg and Peterson, 2009). To determine the phylogenetic relationship between the DST1 copies, the internal element sequence of the DST1 elements was first trimmed with trimal (Capella-Gutiérrez et al., 2009) with parameter ‘-automated1’. Trimmed internal sequences were aligned with MUSCLE (Edgar, 2004) using default settings with manual inspection and a tree was generated with RaxML (Stamatakis, 2014) with settings ‘-m GTRGAMMA -p 12345 -x 12345 -# autoMRE’. To examine expression of individual DST1 elements, diagnostic SNPs were identified for each member of the DST1 family from the multiple sequence alignment and expression per individual element was quantified by requiring a minimum of four RNAseq reads supported by a diagnostic SNP for each sample.

### Annotation of TEs

The ME034V repetitive elements were previously identified using a homology-based repeat masking approach (Thielen *et al*., 2020). We were interested in monitoring potentially active TEs that have intact structural elements, and therefore performed a TE annotation using the EDTA software (Ou *et al*., 2019). This approach was implemented using EDTA v1.9.6 with “--species others --sensitive 1 --anno 1” and all remaining parameters as default. This produced an initial structural annotation of 9,459 intact elements. Simple repeats (‘target_site_duplication, ‘repeat_region’, ‘long_terminal_repeat’) were removed, resulting in a filtered structural annotation of 6,369 intact elements accounting for 16.9 Mb (referred to as “structural annotation”). These structural elements were then used for a homology search, which identified an additional 188,707 elements (115 Mb) that have similarity to structural TEs, but lack intact structural features.

## Data summary statement

The sequences used to profile DNA methylation (EM-Seq) and gene expression (RNA-seq) are available at the National Center for Biotechnology Information (NCBI) BioProject PRJNA787965. Transposable element annotations used in this manuscript are available at https://hdl.handle.net/11299/225624.

## Acknowledgements

Peter Hermanson contributed to the molecular analyses that were important for this study. Michelle Stitzer provided helpful comments and discussion regarding the DST1 TE family. The Minnesota Supercomputing Institute at the University of Minnesota provided computational resources that contributed to this research. This work was funded by NSF IOS-1934384 to N.M.S. and C.N.H. A.R. is supported by NSF PRFB IOS-2109697.

---

**Figure S1. DRM genes in *Setaria viridis*.** (A) The physical location of *DRM* genes in the Setaria viridis ME034V genome is shown. *DRM1a* and *DRM1b* encode putatively functional *DRM* genes and are located in linked positions on chromosome 7. A *DRM3*-like gene on chromosome 3 has mutations in the catalytic domain predicted to disrupt function. (B) The relative expression (tags per million - TPM) of the three *DRM* genes was determined in seedling leaf tissue for wild-type plants, plants derived from tissue culture and the double mutant *drm1ab*. (C) The relative expression of *DRM1b* or *DRM3* compared to *DRM1a* in leaf tissue from several experiments is shown. Dashed line indicates expression level equal to that of *DRM1a*. In most experiments, the expression level of *DRM1a* is much higher than the other two genes. A10 Transcriptome data is from Thielen *et al*. 2020. ME034V data is from this study and Mamidi *et al*., 2020.

**Figure S2. Genome editing reagents and genotyping data.** (A) map of the T-DNA used in transformation to generate edited ME034V. The T-DNA encodes Hygromycin resistance, a wheat codon optimized Cas9 protein encoded as a polyprotein with Trex2 followed by a P2A protein cleavage site, an array of guide RNAs separated by tRNA cleavage sites, and a luciferase reporter. (B) Sanger sequencing results from transgene negative edited *drm1ab* plants.

**Figure S3. Comparisons of DNA methylation levels in drm1ab and unedited plants.** (A) Genome-wide DNA methylation levels for a single replicate of wild-type as well as each of the three replicates of tissue-culture derived and drm1ab plants. (B) Each 100bp tile (n=3,968,817) of the ME034V genome was classified based on the relative levels of mCG, mCHG and mCHH as described in Crisp et al., 2020. The proportion of tiles classified as mCG only, mCG and mCHG, mCHH, intermediate, missing/no data, or unmethylated. (C) The number of tiles classified as mCHH in wild-type, tissue-culture and drm1ab plants. (D) 100bp genomic tiles were classified as ‘high’ (greater than 20%), ‘moderate’ (between 5 and 20%) and ‘low’ (less than 5%) mCHH. A histogram of the percent mCHH methylation lost in drm1ab relative to tissue-culture control plants is shown for each subset of tiles.

**Figure S4. Methylation changes in the drm1ab edited line.** Number of hyper- and hypo-methylated mCG, mCHG, and mCHH tiles when comparing tiles from Tissue Culture to *drm1ab*.

**Figure S5. DRM-dependent loss of mCG and mCHG at regions demarcating edges between high and low mC levels.** A snapshot of a genomic region with a near complete loss of mCHH accompanied by a reduction of mCHG and mCG at, and near, the locations of drm-dependent mCHH.

**Figure S6. A principal component analysis was used to compare expression profiles of drm1ab mutant with wild-type and tissue culture controls.** The drm1ab edited plants form a unique cluster distinct from unedited plants, regardless of whether the unedited plants have been through the tissue-culture process.

