## Supplementary figures and images for "Limited consequences for loss of RNA-directed DNA methylation in *Setaria viridis* domains rearranged methyltransferase (DRM) mutants"

### Supplemental Figure 1

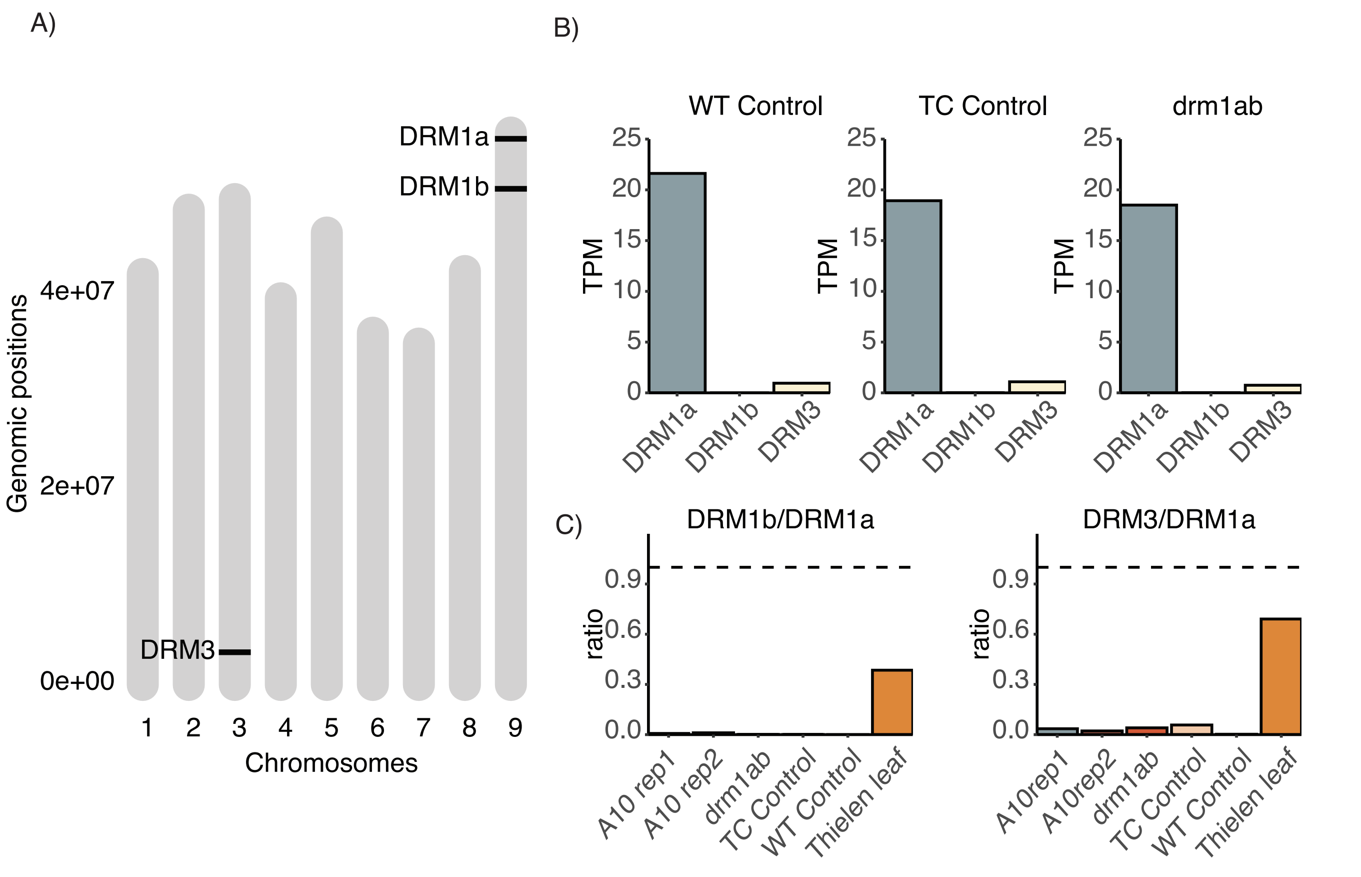

### Supplemental Figure 2

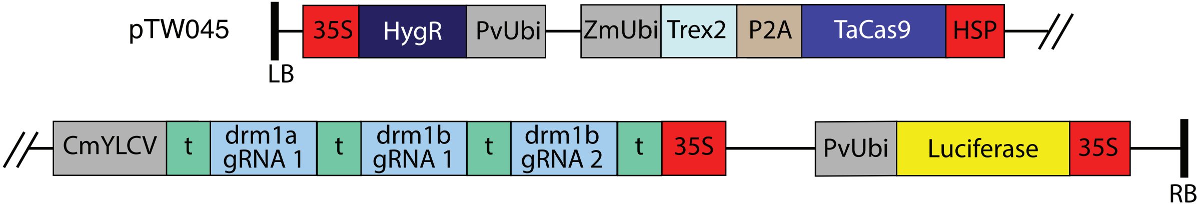

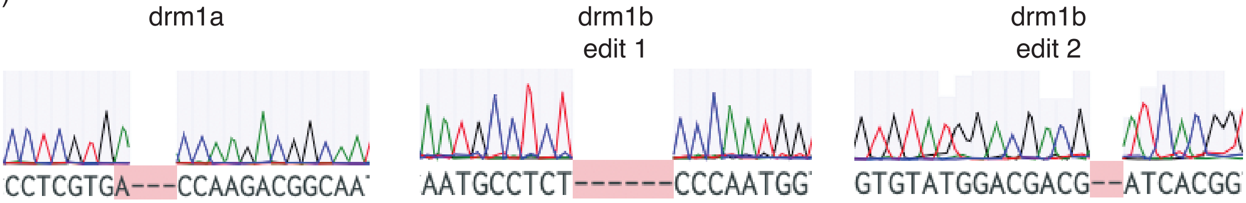

### Supplemental Figure 3

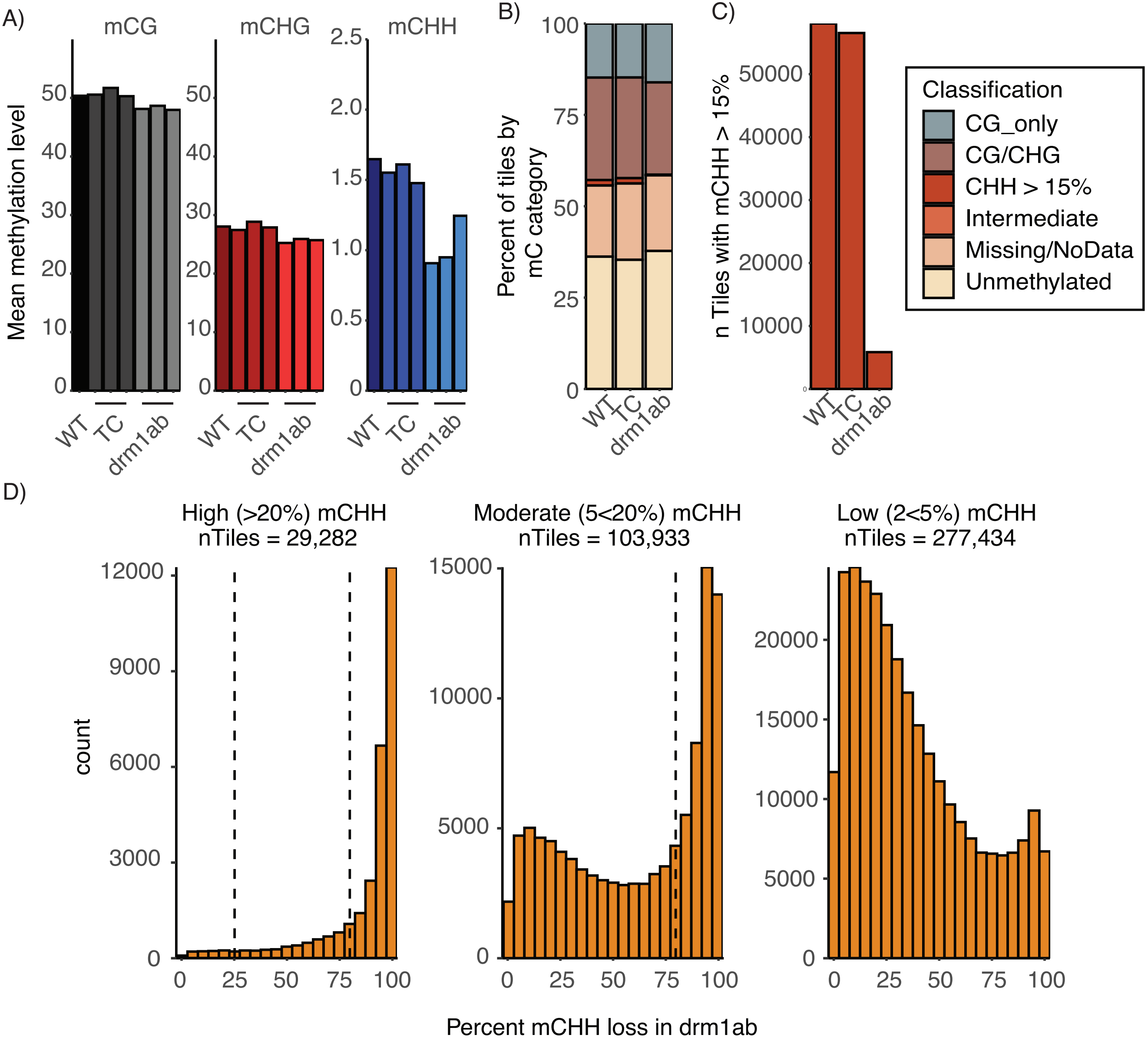

### Supplemental Figure 4

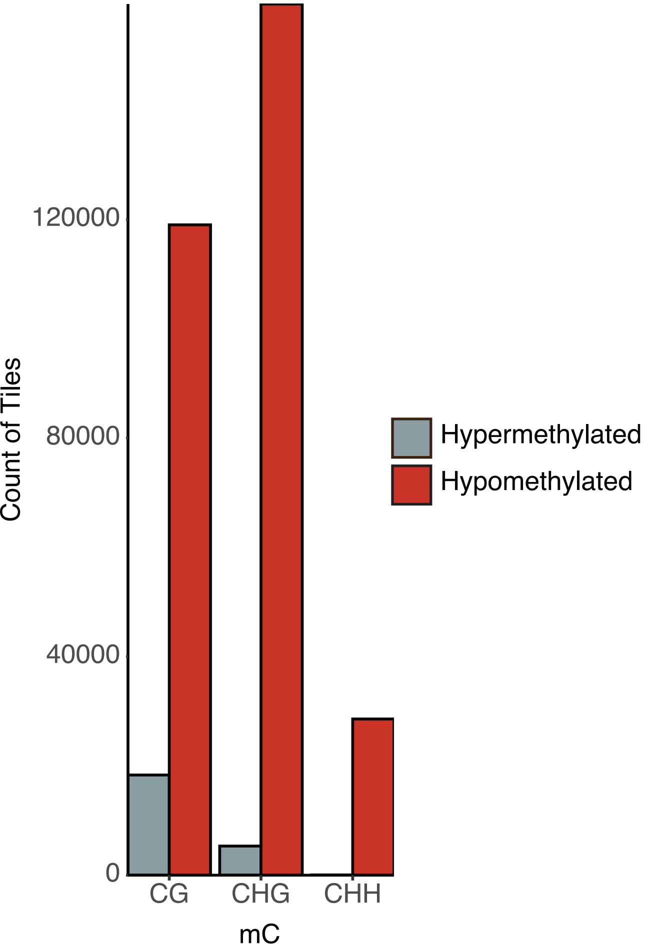

### Supplemental Figure 5

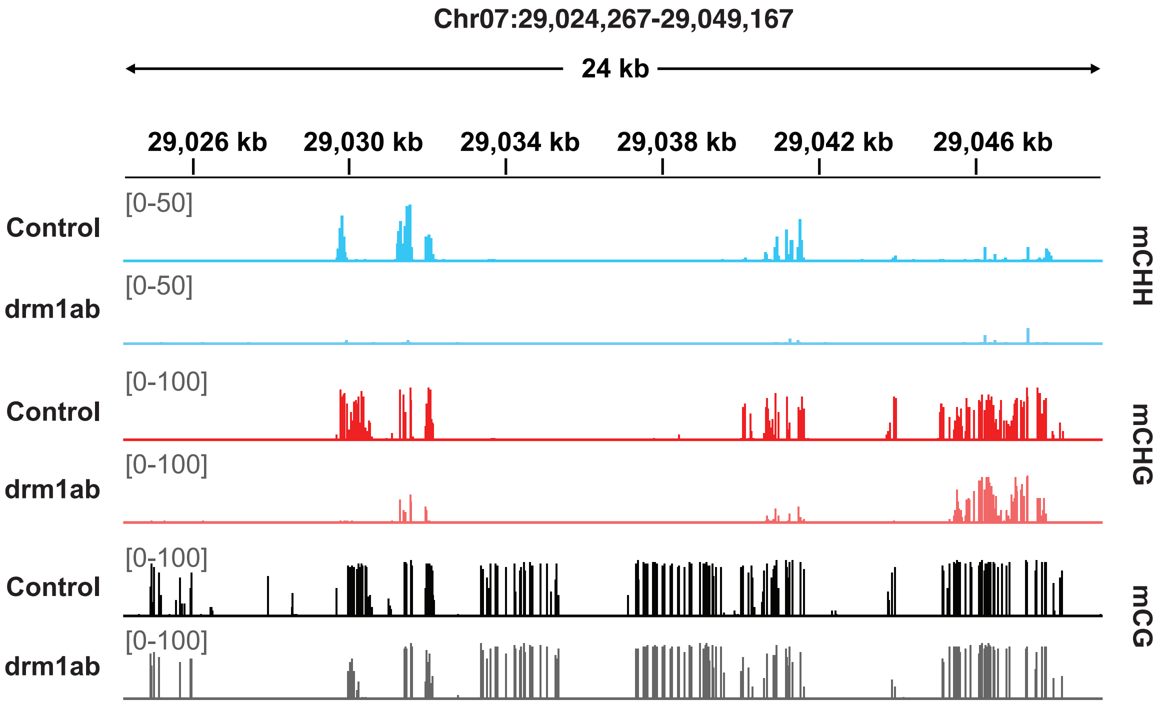

### Supplemental Figure 6

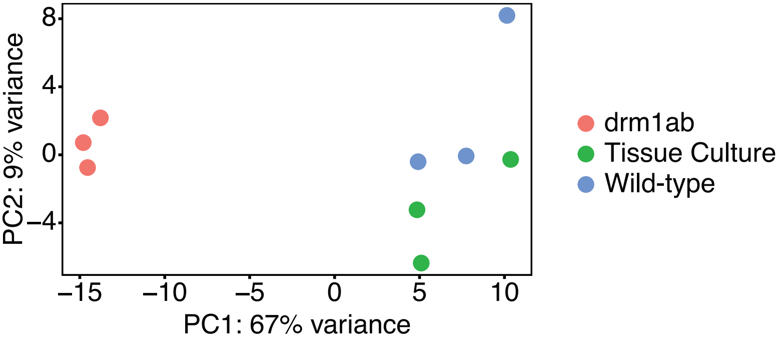
